# Correcting for batch effects in case-control microbiome studies

**DOI:** 10.1101/165910

**Authors:** Sean M. Gibbons, Claire Duvallet, Eric J. Alm

**Affiliations:** Department of Biological Engineering, Massachusetts Institute of Technology, Cambridge, MA, USA; Center for Microbiome Informatics and Therapeutics, Cambridge, MA, USA; The Broad Institute of MIT and Harvard, Cambridge, MA, USA

## Abstract

High-throughput data generation platforms, like mass-spectrometry, microarrays, and second-generation sequencing are susceptible to batch effects due to run-to-run variation in reagents, equipment, protocols, or personnel. Currently, batch correction methods are not commonly applied to microbiome sequencing datasets. In this paper, we compare different batch-correction methods applied to microbiome case-control studies. We introduce a model-free normalization procedure where features (i.e. bacterial taxa) in case samples are converted to percentiles of the equivalent features in control samples within a study prior to pooling data across studies. We look at how this percentile-normalization method compares to traditional meta-analysis methods for combining independent p-values and to limma and ComBat, widely used batch-correction models developed for RNA microarray data. Overall, we show that percentile-normalization is a simple, non-parametric approach for correcting batch effects and improving sensitivity in case-control meta-analyses.

**Author Summary:** Batch effects are obstacles to comparing results across studies. Traditional meta-analysis techniques for combining p-values from independent studies, like Fisher’s method, are effective but statistically conservative. If batch-effects can be corrected, then statistical tests can be performed on data pooled across studies, increasing sensitivity to detect differences between treatment groups. Here, we show how a simple, model-free approach corrects for batch effects in case-control microbiome datasets.

## Introduction

Data generated by high throughput methods like mass-spectrometry, second-generation sequencing, or microarrays are sensitive to experimental and computational processing [1, 2]. This sensitivity gives rise to ‘batch effects’ between independent runs of an experiment. Even when different research groups adhere to the same methodologies, these effects can arise due to slight differences in hardware, reagents, or personnel [3]. Thus, it is inappropriate to make direct, quantitative comparisons of uncorrected data across studies.

Several tools for reducing batch effects in RNA microarray data have been developed. For example, surrogate variable analysis (SVA) estimates a set of inferred variables (eigenvectors) that explain variance associated with putative batch effects [4]. These inferred variables are then incorporated into a linear model to correct downstream significance tests. The limma package employs a similar linear correction to account for batch effects prior to statistical analysis [5]. SVA and limma are part of a family of linear batch-correction methods that use different varieties of factor analysis, singular value decomposition, or regression [4–7]. The most relied upon method to date [8], called ComBat, uses a Bayesian approach to estimate location and scale parameters for each feature within a batch [9]. All of these models are most effective when batch effects are not conflated with the true biological effects [1]. Furthermore, most batch correction methods make certain parametric assumptions.

Unfortunately, models that often work well for many types of ‘omics data may not be appropriate for microbiome datasets. In microbiome studies, batch effects are often diffuse and conflated with biological signals [10–12]. The microbiome field has also struggled with finding appropriate parametric models for bacterial abundance distributions and for dealing with zeros. This is especially true for low-biomass samples in microbiome sequencing studies, like samples taken from the built environment [13], where populations are under-sampled, the biological signal is relatively weak, and batch effects can be quite large [14]. One way to get around this issue is to calculate statistics within a given batch, and then compare significant features across batches using classic meta-analysis techniques for combining p-values, like Fisher’s and Stouffer’s methods [15, 16]. These meta-analysis techniques are robust to batch effects across independent studies, but have less statistical power and ability to detect subtle differences than directly pooling data across studies.

Here, we describe a model-free data-normalization procedure for controlling batch effects in case-control microbiome studies that enables pooling data across studies. Case-control studies include a built-in population of control samples (e.g. healthy subjects) that can be used to normalize the case samples (e.g. diseased subjects). For every feature (i.e. bacterial taxon), the case abundance distributions can be converted to percentiles of the equivalent control abundance distributions (Fig. 1). Study-specific batch effects present in the case samples will also be present in the control samples, and by converting the case data into percentiles of the control distribution these effects are mitigated. Upon conversion to percentiles of the within-study controls, percentile-normalized samples from multiple studies with similar case-control definitions can be more appropriately pooled for statistical testing (Fig. 1). We show that this approach effectively controls batch effects in microbiome case-control studies and we compare this method to pooling ComBat- or limma-corrected data, and to Fisher’s and Stouffer’s methods for combining independent p-values.

**Figure 1.**
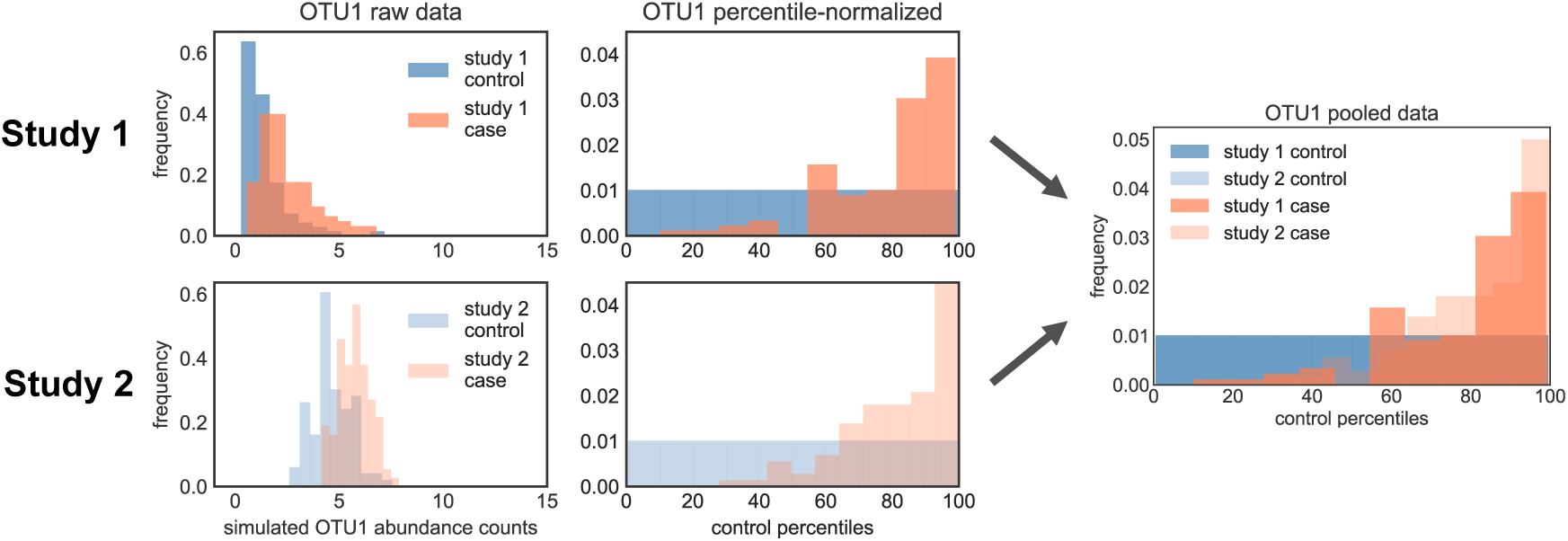
Percentile-normalization procedure converts case and control values into percentiles of the control distribution, which allows for pooling of normalized data across studies. Conceptual plot shows theoretical feature (OTU 1) abundance distributions for control samples and case samples from two independent studies. Converting a control distribution into percentiles of itself naturally gives rise to a uniform distribution (represented by flat blue distributions in central panels), while converting the case distribution into percentiles of the control distribution produces a non-uniform distribution when these two distributions differ (represented by skewed orange distributions in central panels). The right-most panel shows the result of pooling percentile distributions from study 1 and study 2. Percentile-normalization places data from separate studies onto a standardized axis that allows for cross-study comparison. Each simulated case and control distribution was produced by randomly sampling 100 times from a lognormal distribution. Study 1 control parameters: µ = 0.1 and σ = 0.7. Study 1 case parameters: µ = 0.8 and σ = 0.5. Study 2 control parameters: µ = 1.5 and σ = 0.2. Study 2 case parameters: µ = 1.75 and σ = 0.13.

## Methods

### Datasets

We used a collection of case-control datasets obtained from the MicrobiomeHD database [17] to validate our batch-normalization method. We focused our analyses on studies spanning five diseases: colorectal cancer (CRC) [18–21], Crohn’s Disease (CD) [22–25], Ulcerative Colitis (UC) [23–25], obesity (OB) [18, 26–35], and *Clostridium difficile* induced diarrhea (CDI) [33, 36]. For a subset of three CRC studies [18–20], we were able to obtain sequence data from the same region of the 16S gene (V4) so that these data could be processed together. The remaining MicrobiomeHD case-control datasets were previously processed using the same pipeline (see below), and then Operational Taxonomic Units (OTUs) were summarized at the genus level for comparison across studies.

### Sequence Data Processing

To perform OTU-level analyses across the CRC studies, we downloaded the raw data from all of the MicrobiomeHD datasets that sequenced the V4 region of the 16S gene. We quality filtered and length trimmed each V4 dataset as described in Duvallet et al. (2017) and concatenated these raw, trimmed FASTQ files into one file. We removed any unique sequences that did not appear more than 20 times and clustered the remaining reads with USEARCH [37] at 97% similarity. We assigned these OTUs taxonomic identifiers using the RDP classifier [38] with a cutoff of 0.5.

For genus-level analyses, OTU tables and metadata were acquired from the MicrobiomeHD database (https://doi.org/10.5281/zenodo.569601). Raw data were downloaded from the original studies and processed through our in-house 16S-processing pipeline (https://github.com/thomasgurry/amplicon_sequencing_pipeline) as described previously [17]. Each study’s OTU table was converted to relative abundance by dividing each sample by its total number of reads and collapsed to genus level by summing all OTUs with the same genus-level annotation.

To plot data in ordination space, Bray-Curtis distances were calculated from relative abundance data using Scikit-learn (sklearn.metrics.pairwise.pairwise_distances; metric=’braycurtis’) [39]. Non-metric multidimensional scaling (NMDS) coordinates were calculated for two axes based on Bray-Curtis distances using Scikit-learn (sklearn.manifold.MDS; n_components=2, metric=False, max_iter=500, eps=1e-12, dissimilarity=’precomputed’).

### Percentile Normalization

In this procedure, control feature distributions are percentile-normalized against themselves (resulting in a uniform distribution between 0 and 100) and case feature distributions are converted into percentiles of their equivalent control features. Treating our controls as null-hypotheses is motivated by the idea that healthy patients should be treated as similar across datasets, even though we understand that they will differ due to biological as well as technical batch effects. Relative abundance distributions were converted to percentiles using the SciPy v 0.19.0 [40] stats.percentileofscore method (kind=’mean’). In order to avoid rank pile-ups due to the presence of many zeros, we replaced zeros with pseudo relative abundances drawn from a uniform distribution between 0.0 and 10^-9^ (i.e. a set of random values smaller than the lowest possible relative abundance in any dataset). Due to the zero-replacement step, p-values can shift slightly upon re-analysis with a different random draw, which can lead to the loss or gain of significance for features very near the significance threshold. Within each study, control distributions for each individual OTU or genus were converted into percentiles of themselves and case distributions were converted into percentiles of their corresponding control distribution. We have written a python script that performs percentile-normalization given an OTU table, a list of case sample IDs, and a list of control sample IDs as inputs (https://github.com/seangibbons/percentile_normalization). A QIIME 2 (https://qiime2.org) plugin for running percentile-normalization is also available (https://github.com/cduvallet/q2-perc-norm).

### ComBat

For each disease, we applied ComBat [8] to the case-control datasets analyzed in this study. Relative abundances (OTUs in the CRC analysis or OTUs collapsed to the genus level in the genus-level analysis) were log-transformed prior to running ComBat (default settings), adding a pseudo relative abundance of half the minimal frequency (across the entire feature table) to replace zeros. ComBat-corrected data were then transformed back from log-space (i.e. exponential transformation) prior to downstream analyses.

### limma

In addition to ComBat, we applied a linear batch correction method from the limma package in R [5]. Relative abundances (zeros replaced with pseudo relative abundances equal to half the minimal frequency across the entire feature table) were log-transformed as described above and then a linear model was fit to subtract batch effects using the removeBatchEffect function (default settings). The limma-corrected data were then transformed back from log-space (i.e. exponential transformation) prior to downstream analysis.

### Statistical Analysis

To calculate statistical significance, we restricted our statistical tests to OTUs/genera that occurred in at least one third of control or one third of case samples in order to reduce our multi-test correction penalty. We used the Wilcoxon rank-sum test, as implemented in SciPy v0.19.0 (sicipy.stats.ranksums) [40], to determine significant differences between independent groups of samples. Wilcoxon tests were run either within or across studies. In order to calculate statistics across studies, normalized case and control samples from multiple studies of the same disease were combined together into the same OTU table. Hereafter, combining datasets is referred to as ‘pooling.’ P-values were multiple-test corrected using the Benjamini-Hochberg False Discovery Rate (FDR) procedure, as implemented in StatsModels v 0.8.0 (statsmodels.sandbox.stats.multicomp.multipletests) [41]. Differences in overall community structure were assessed using the Permutational Multivariate Analysis of Variance (PERMANOVA) test in R’s vegan package [42] as implemented in scikit-bio (skbio.stats.distance.permanova). Fisher’s and Stouffer’s methods for combining p-values were performed using SciPy v0.19.0 (scipy.stats.combine_pvalues; method=’fisher’ or method=’stouffer’). For Stouffer’s method, weights for each study were calculated as the square root of the number of cases plus the number of controls. OTUs/genera with significant responses in opposing directions across studies were excluded from Fisher and Stouffer analyses.

### in silico experiments

We ran an *in silico* titration experiment using the OTU-level data to simulate pooling of control samples from different datasets before calculating significant differences. Healthy samples from one study were mixed with healthy samples from another study at different proportions prior to calculating significant differences in OTU frequencies between cases and controls. Case and control groups were subsampled to 40 samples each. Control samples were substituted by randomly selected samples from another study along a fractional gradient (0-100% control samples from another study). We calculated significant differences between case and control groups using the Wilcoxon rank-sum test and applied an FDR correction. OTUs with q-values ≤ 0.05 were considered significant. The titration experiment was rerun 20 times, and the results were averaged.

Similar to the titration experiment, we ran an OTU-level analysis of how batch-correction methods might impact false-positive rates by randomly selecting 40 control samples from the Baxter et al. (2016) study as artificial ‘controls’ and 40 control samples from the Zeller et al. (2014) study as artificial ‘cases’ (across 20 iterations) for each data type (i.e. raw, percentile-normalized, limma-corrected, and ComBat-corrected). We then calculated significant differences between these artificial ‘case’ and ‘control’ groups as outlined above to generate p-values for each OTU.

## Results

### Batch effects at OTU-level resolution

To minimize possible biases across data sets, we identified three colorectal cancer (CRC) studies that sequenced the same region of the 16S gene (V4). We reprocessed the raw sequence data from each study in the same quality filtering and OTU picking pipeline to obtain bioinformatically-standardized results. OTUs that occurred in at least one third of case or one third of control samples (i.e. within individual studies) were retained for all downstream statistical analyses. Despite standardizing the bioinformatic processing of these data, we saw significant batch effects in healthy patients across studies (PERMANOVA p < 0.001; Fig. 2). The similarity between samples from the Baxter et al. (2016) and Zackular et al. (2014) studies is due to the fact that they were sourced from the same patient cohort (although samples were processed separately), making this comparison a good pseudo-negative control for batch effects [18, 20]. There was an apparent reduction in the batch effect after applying ComBat, although differences between batches remained weakly significant (PERMANOVA p = 0.008, Fig. 2) [8]. Due to the non-independence between the Baxter and Zackular patient cohorts, we removed the smaller of the two studies (Zackular) from all downstream analyses. Out of a total of 1,021 OTUs that passed our abundance filter, 681 differed significantly in uncorrected relative abundance between the Baxter and Zeller healthy controls (FDR q ≤ 0.05).

**Figure 2.**
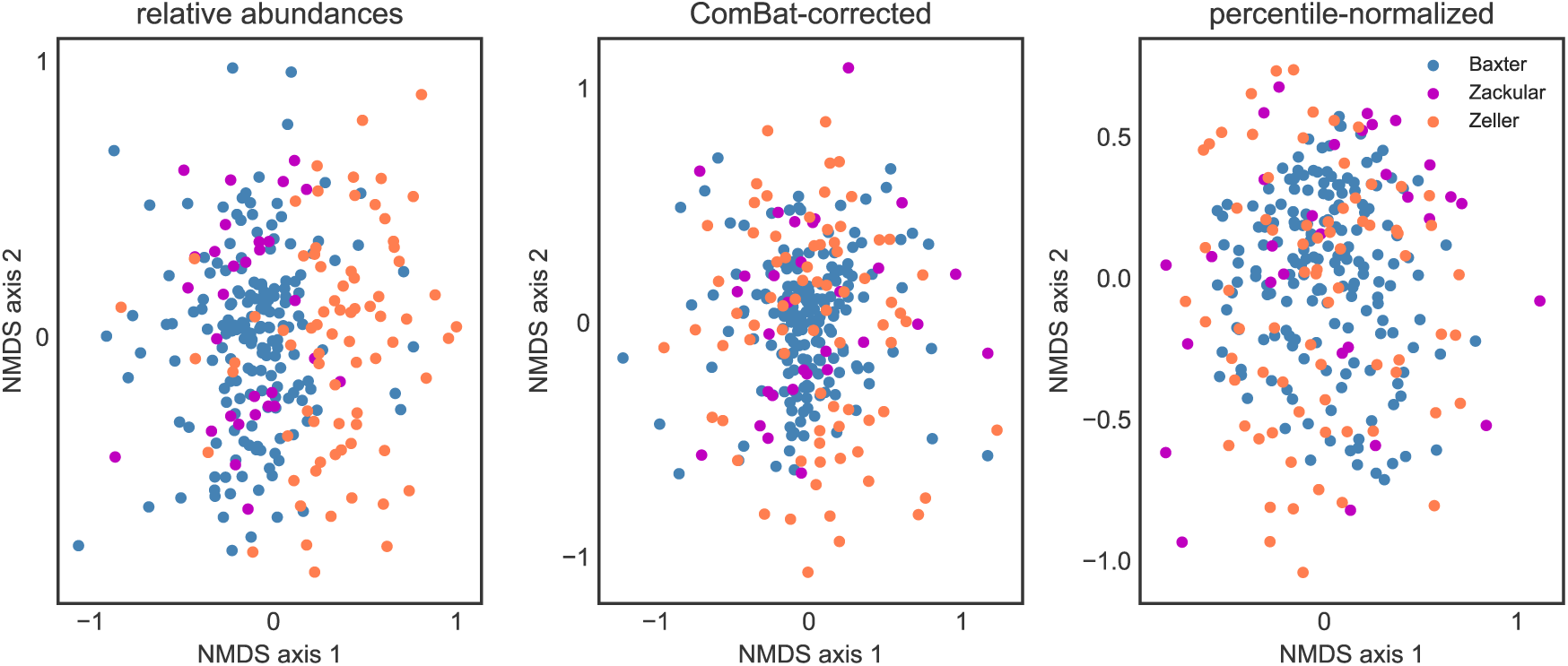
Batch effects between healthy controls from different studies can be reduced by ComBat and percentile-normalization. Non-metric multidimensional scaling (NMDS) plot showing the distribution of healthy controls from three colorectal cancer studies in ordination space (Bray-Curtis distances of relative abundance OTU-level data). Despite standardized bioinformatic processing, healthy patients differed significantly in their gut microbiomes across studies (PERMANOVA p < 0.001; batch accounts for 6.342% of the total variance). Studies were still significantly different after applying ComBat, an established batch-correction method (PERMANOVA p < 0.01). However, percentile-normalization did a better job of stabilizing the variance across studies and removed any apparent batch effect (PERMANOVA p > 0.5).

We ran an *in silico* titration experiment to simulate pooling of control samples from different datasets before calculating significant differences. Healthy samples from one study were mixed with healthy samples from another study at different proportions prior to calculating significant differences in OTU frequencies between cases and controls (see conceptual outline in Fig. 3). For non-normalized data, the number of significant OTUs greatly increased due to batch effects as more control samples were substituted in from another study. This result highlights the danger of pooling raw data across batches. ComBat- and limma-corrected data performed better than uncorrected data, but still showed many spurious results as the proportion of control samples from another study increased (Fig. 3). Percentile-normalization showed no increase in spurious results over the titration gradient (Fig. 3). Although we do see several significant CRC-associated OTUs in the full dataset (see below), these were not detected in the titration experiment due to the loss of statistical power when reducing case and control groups to only 40 samples each.

**Figure 3.**
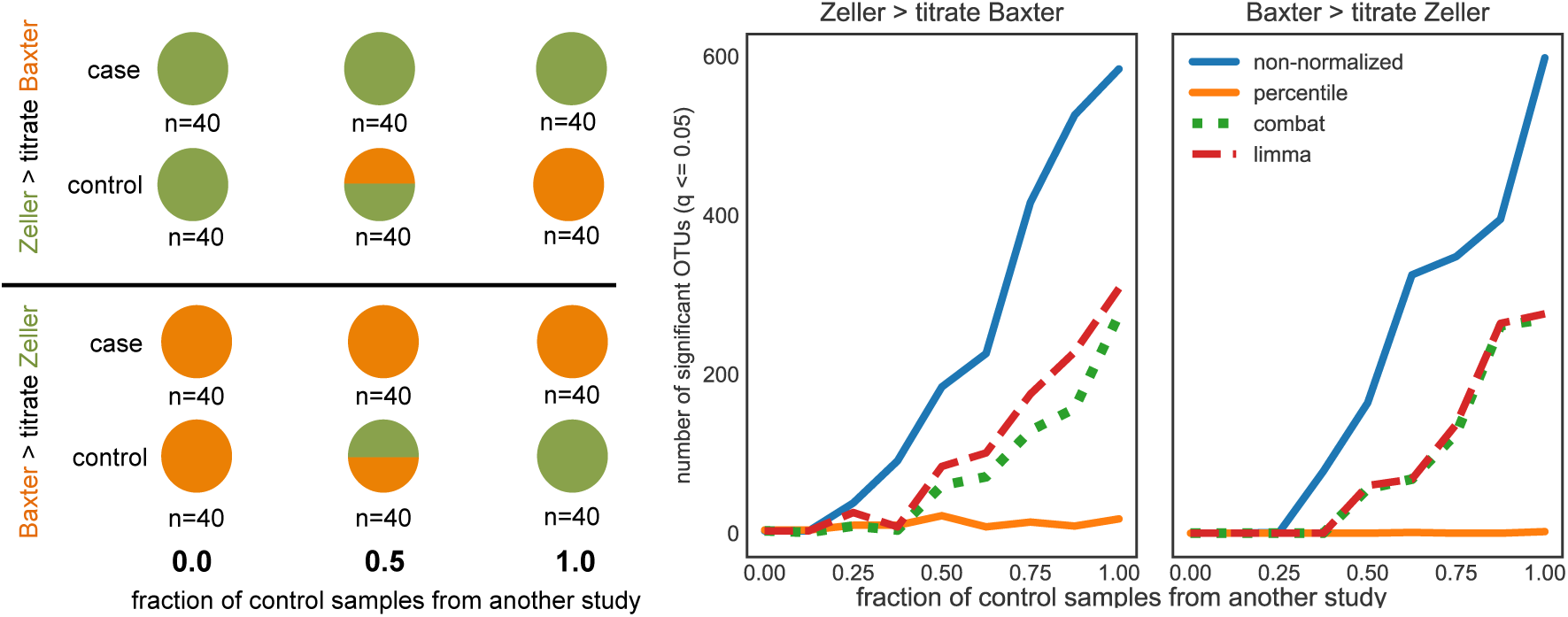
Pooling non-normalized samples from different studies can give rise to many spurious associations. The control group from one study is gradually substituted with randomly chosen control samples from another study (non-normalized, percentile-normalized, limma-corrected, and ComBat-corrected), keeping the total number of case and control samples fixed at n=40 (see conceptual illustration on the left). Mixing in non-normalized control samples from another study gave rise to spurious results due to batch effects (blue lines). ComBat- and limma-corrected data showed fewer spurious associations (green and red lines). Percentile-normalization showed no increase in spurious results along the titration gradient (orange lines).

We ran a second *in silico* experiment to determine whether the false-positive rate was impacted by our different batch-correction methods. We randomly selected 40 samples from the Baxter healthy controls as artificial ‘controls’ and 40 samples from the Zeller healthy controls as artificial ‘cases’ for each data type (i.e. non-normalized, percentile-normalized, limma-corrected, and ComBat-corrected) and calculated significant OTU-level differences between these groups. We repeated this process twenty times to generate a set of p-value distributions. We found that the fraction of p-values ≤ 0.05 can be as high as ∼70% for the non-normalized data (Fig. 4). This result matches with our finding that 681 out of the 1,021 OTUs in this dataset differed significantly across Baxter and Zeller controls (q ≤ 0.05). Each normalization technique drastically reduced the number of false positives, but percentile-normalization gave the best results (Fig. 4). When low-abundance OTUs were included in the analysis, ComBat and limma showed highly skewed p-value distributions, giving rise to a larger number of false positives than the non-normalized data (Fig. S1).

**Figure 4.**
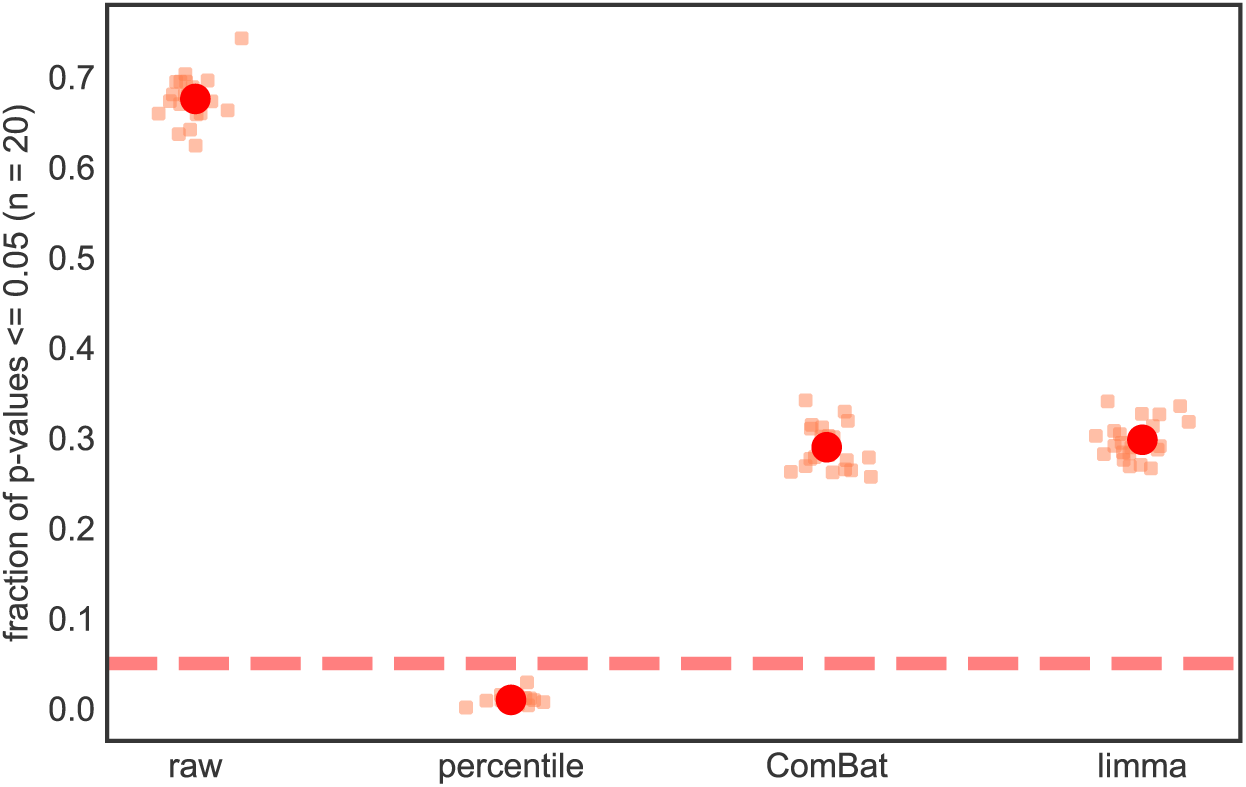
False positive rates are reduced by batch-correction methods. Random sets of 40 Baxter controls and random sets of 40 Zeller controls were selected for null case-control comparisons (20 iterations). Smaller points show the fraction of p-values ≤ 0.05 within a given iteration, while larger dots show the average value across all 20 iterations. Within each category, smaller points are randomly jittered along the x-axis for better visualization. The fraction of p-values ≤ 0.05 is highly inflated for non-normalized data (red dashed line shows the null-expectation for p-values). Only abundant OTUs (detected in at least a third of case or control samples) were included in this analysis.

We next assessed the performance each cross-study analysis method by comparing OTU-level results across two independent CRC datasets. In the Baxter study, there were 172 healthy (control) samples and 120 CRC (case) samples, with 14 OTUs (from *Fusobacterium*, *Coprococcus*, *Butyricicoccus*, *Gemmiger*, *Faecalibacterium*, *Roseburia*, *Parvimonas*, *Haemophilus*, *Porphyromonas*, *Peptostreptococcus*, *Streptophyta*, *Bacteroides* and *Clostridium XIVa* genera) showing significant differences in abundance between cases and controls (FDR q ≤ 0.05). For Zeller, there were 71 control and 40 case samples, with 18 OTUs (from *Butyricicoccus*, *Butyricimonas*, *Fusobacterium*, *Closridium XIVa*, *Streptococcus*, *Parabacteroides*, *Alistipes*, *Anaerostipes*, *Parvimonas*, *Peptostreptococcus*, *Blautia*, *Dialister,* and *Bacteroides* genera) that differed significantly across cases and controls (FDR q ≤ 0.05).

In the absence of batch effects, pooling data across datasets of the same disease should increase sensitivity to detect significant cross-study associations. We pooled percentile-normalized, limma-corrected, and ComBat-corrected data, respectively, across Baxter and Zeller studies to look for OTUs that differed significantly across cases and controls. These pooled results were then compared to classic methods for combining p-values from each dataset’s individual results (above). For the percentile-normalized data, we found 39 OTUs that differed significantly between cases and controls (FDR q ≤ 0.05), 21 of which overlapped with the within-study results. The pooled limma-corrected and ComBat-corrected data resulted in 37 and 36 significant OTUs, respectively. 35 of the OTUs identified as significant by ComBat were also significant in the limma results. 30 of the limma results and 29 of the ComBat results were also significant in the percentile-normalization results, respectively. Fisher’s method identified seven significant OTUs from *Clostridium XIVa*, *Streptococcus*, *Fusobacterium*, *Parvimonas*, *Peptostreptococcus*, and *Anaerostipes* genera, which were also found in the percentile-normalized results. Stouffer’s method identified the same seven OTUs found using Fisher’s method. Overall, the pooling methods improve statistical power to detect significant OTUs over traditional meta-analysis methods. For example, particular OTUs from *Desulfovibrio* and *Parabacteroides* genera were identified as significantly enriched in CRC patients in the pooled results (ComBat, limma, and percentile-normalized), but not in the within-study results or in the Fisher and Stouffer results. Pooled analysis of percentile-normalized data also identified on *Enterobacter* OTU enriched in cancer patients, two OTUs from the *Lachnospiraceae* family that were enriched in controls and one *Lachnospiraceae* that was enriched in cases, which were missed by the within-study analyses. In all, 18 OTUs were identified in the pooled, percentile-normalized results that were missed by the within-study analyses (Fig. 5). These additional taxonomic associations (e.g. *Desulfovibrio*, *Costridium XIVa*, and *Lachnospiraceae*) are consistent with prior meta-analyses of CRC microbiome studies [17, 43]. It is important to visualize the data being fed into statistical tests to determine whether significant associations are being driven by outlier studies or by other artifacts. The associations identified in Figure 5 appear to be biologically meaningful due to the overall consistency of the effect directions across studies.

**Figure 5.**
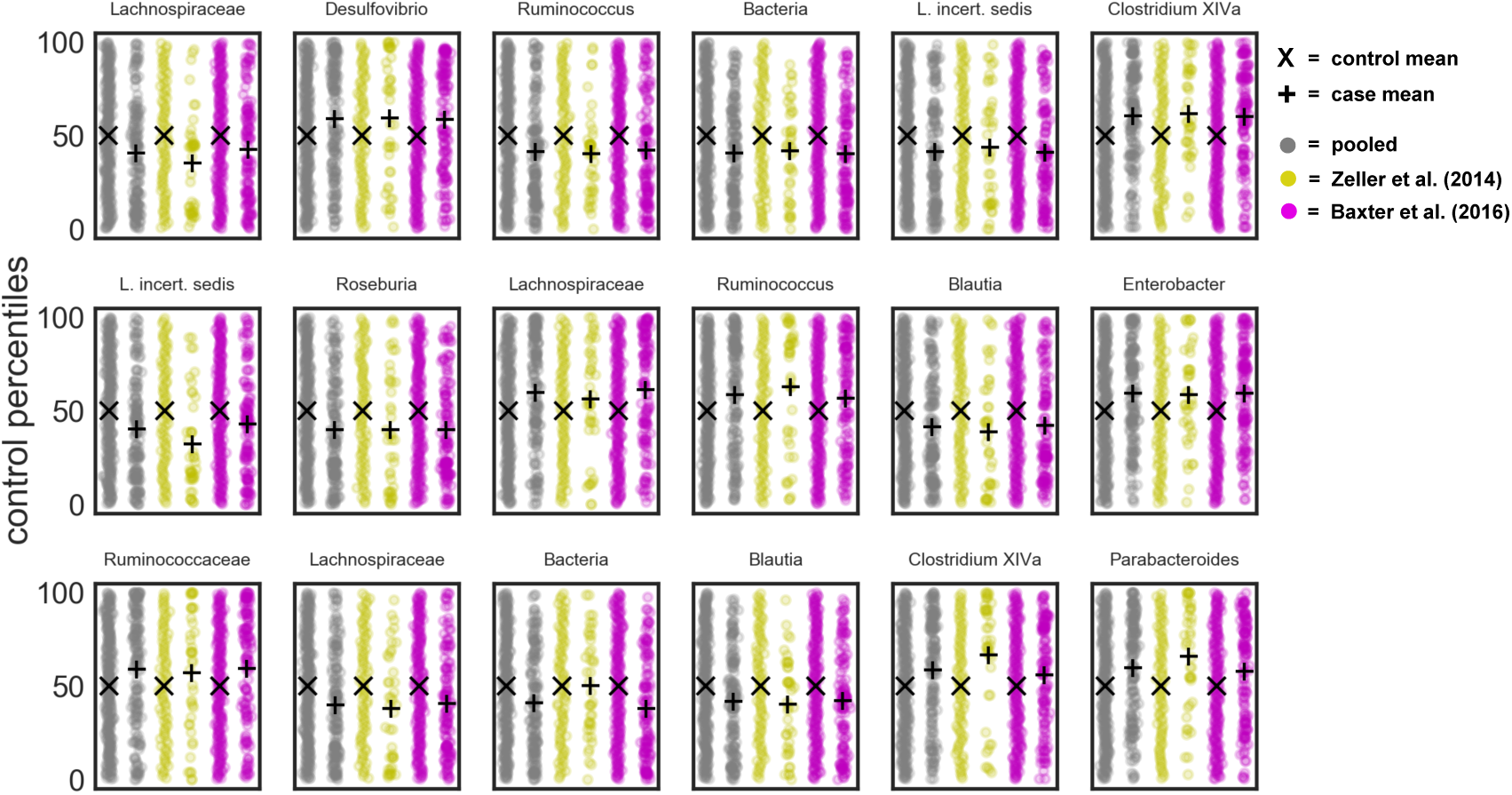
OTUs significant across CRC studies, but not within a given study. Pooling data provides greater statistical power to detect subtle, yet consistent differences in OTU abundances across sample groups. 18 OTUs are labeled by their most resolved taxonomic annotation. Each OTU in this plot was not found to be significant within either Baxter or Zeller studies, but became significant after pooling the percentile-normalized datasets (q ≤ 0.05).

### Batch effects at genus-level resolution across multiple diseases

In order to assess the performance of different meta-analysis techniques across a larger set of studies and diseases, we summarized OTU abundances at the genus level for five diseases across 18 studies - *Clostridium difficile induced diarrhea* (CDI), Crohn’s disease (CD), ulcerative colitis (UC), obesity (OB), and CRC. There were a total of 306 unique genera detected across studies. There were two CDI case-control studies: Schubert et al. (2014) had 154 control and 93 case samples [33]; Vincent et al. (2013) had 25 control and 25 case samples [36]. There were four inflammatory bowel disease (IBD) studies that included CD patients and three that also included UC patients: Papa et al. (2012) had 24 non-IBD control samples, 23 CD samples, and 43 UC samples [23]; Morgan et al. (2012) had 18 control, 61 CD and 47 UC samples [24]; Willing et al. (2010) had 35 control, 16 UC and 29 CD samples [25]; Gevers et al. (2014) had 16 non-IBD control and 146 CD samples, with no UC samples [22]. There were eleven studies with lean and obese (OB) cohorts: Turnbaugh et al. (2009) had 33 controls and 102 cases [26]; Goodrich et al. (2014) had 428 controls and 185 cases [30]; Escobar et al. (2014) had 10 controls and 10 cases [27]; Zhu et al. (2014) had 16 controls and 25 cases [44]; Jumpertz et al. (2011) had 12 controls and 9 cases [34]; Ross et al. (2015) had 26 controls and 37 cases [29]; Zupancic et al. (2012) had 96 controls and 101 cases [28]; Baxter et al. (2016) had 125 controls and 47 OB cases [18]; Schubert et al. (2014) had 68 controls and 34 OB cases [33]; Wu et al. (2011) had 59 controls and 9 cases [32]; and Zeevi et al. (2015) had 567 controls and 151 cases [31]. There were four independent CRC studies, including the Baxter and Zeller studies listed in the OTU-level analysis (see above for sample sizes). The remaining two CRC studies are Wang et al. (2012), which had 54 control and 44 case samples [45], and Chen et al. (2012), which had 22 controls and 21 cases [21].

Most genera that differed significantly within a given study were not significant in other studies of the same disease. For example, of the 36 unique genera that showed significant differences within any given OB study, none were found to be significant in all studies (I = 0; Table 1). Indeed, there were no genera that were significant across all studies in the majority of diseases studied (I = 0; Table 1). CDI only had two studies, and of the 38 significant results, only six were shared across both datasets. Overall, few genera were significant within two or more studies (2N ≤ 6; Table 1).

**Table 1.** Normalization methods impact the number of significant genus-level associations between cases and controls across multiple diseases. Numbers of genera that differ significantly between cases and controls for five diseases. In the ‘disease’ column, CDI = *Clostridium difficile* induced diarrhea, CD = Crohn’s Disease, UC = Ulcerative Colitis, CRC = Colorectal Cancer, and OB = obesity. ‘N =’ shows the number of studies included in each meta-analysis. The method column indicates how the data were processed prior to running significance tests (percentile-normalized, Fisher’s method for combining p-values, Stouffer’s method for combining p-values, ComBat-corrected, or limma-corrected). The significance threshold used was q ≤ 0.05 (FDR). The ‘pooled’ column shows the total number of genera that were found to be significantly different *across* pooled studies for a given disease. The ‘within’ column shows the total number of unique (non-redundant) genera that were identified as significantly different *within* each study (U = union), the number of genera that were significant in all individual studies (I = intersection), and the number of significant genera that were consistently significant in at least two studies (2N; 2N == I for CDI).

| disease | method | pooled | within |
| --- | --- | --- | --- |
| CDI (N=2) | percentile | 37 | U = 38 / 2N = 6 |
|  | Fisher | 12 |  |
|  | Stouffer | 12 |  |
|  | ComBat | 36 |  |
|  | limma | 36 |  |
| CD (N = 4) | percentile | 19 | U = 13 / I = 0 / 2N = 2 |
|  | Fisher | 6 |  |
|  | Stouffer | 6 |  |
|  | ComBat | 2 |  |
|  | limma | 1 |  |
| UC (N = 3) | percentile | 10 | U = 17 / I = 0 / 2N = 1 |
|  | Fisher | 4 |  |
|  | Stouffer | 4 |  |
|  | ComBat | 5 |  |
|  | limma | 5 |  |
| CRC (N = 4) | percentile | 12 | U = 20 / I = 0 / 2N = 3 |
|  | Fisher | 9 |  |
|  | Stouffer | 7 |  |
|  | ComBat | 5 |  |
|  | limma | 5 |  |
| OB (N = 11) | percentile | 18 | U = 36 / I = 0 / 2N = 6 |
|  | Fisher | 4 |  |
|  | Stouffer | 6 |  |
|  | ComBat | 13 |  |
|  | limma | 15 |  |

The number of genera that differed significantly across pooled cases and controls changed depending on how the data were batch-corrected (Table 1 and Table S1). For every disease, percentile-normalization yielded the largest number of significant genera when compared to other methods. Overall, ComBat- and limma-corrections resulted in many fewer significant genera, especially for CD, UC, and CRC (Table 1 and Table S1). Half of the IBD (CD and UC) studies included non-IBD patients with inflammatory symptoms as controls rather than clinically healthy patients. These biologically relevant differences in inflammatory symptoms between control cohorts were conflated with batches and were likely smoothed out by ComBat and limma corrections. Fisher’s and Stouffer’s methods consistently identified fewer significant associations than percentile-normalization (Table 1 and Table S1). Pooling data prior to running a statistical test is a more sensitive technique than combining independently calculated p-values [46]. Thus, percentile-normalization increases the statistical power to detect differences across studies while controlling for false positives and batch effects.

To better assess how percentile normalization impacted the pooled results, we looked at genera that were significant within a single-study but not across studies after pooling. There were 12 genera that were significant within a subset of CRC studies, but not after pooling (Fig. 6). *Gemmiger*, *Bacteroides*, and *Roseburia* showed variable responses across studies, sometimes enriched in controls and other times enriched in cases. The remaining genera showed weak associations within one or two studies, but did not differ significantly across studies (i.e. q ≤ 0.05). These genera that show weak or inconsistent responses across batches may not be reliable disease biomarkers. However, by including larger numbers of CRC studies in future meta-analyses it is likely that some of these genera could pass the significance threshold. Two genera - *Lactobacillus* and *Desulfovibrio* - were not significantly different between cases and controls within an individual study, but became significant after pooling (Fig. 7). These genera showed weak, but largely consistent enrichment in cancer patients and demonstrate the utility of pooling datasets to detect subtle differences.

**Figure 6.**
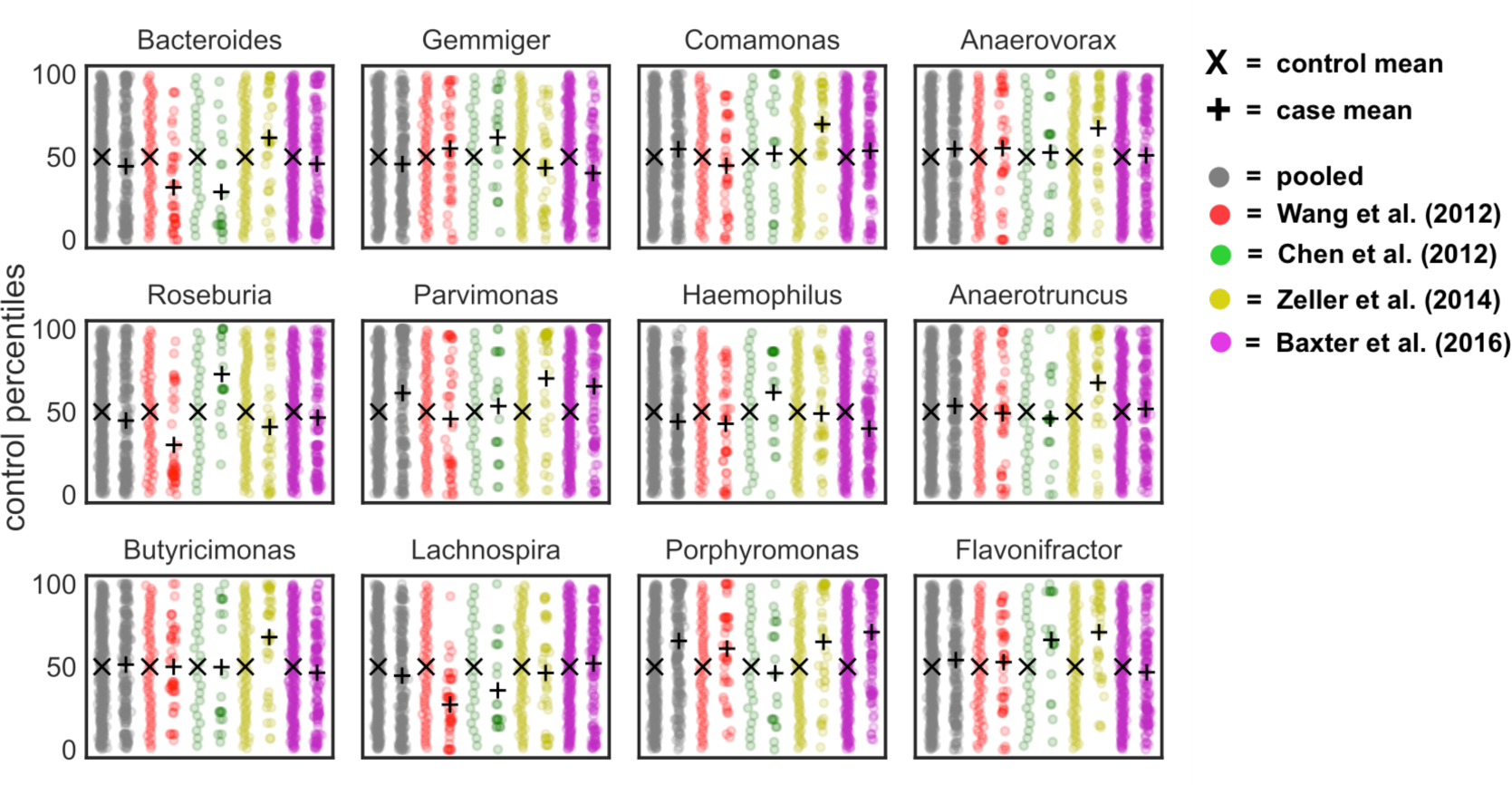
Genera that show a significant difference between CRC cases and controls within a given study, but not after pooling. 12 genera showed significant differences between cases and controls within a study (q ≤ 0.05), but not after pooling across CRC studies.

**Figure 7.**
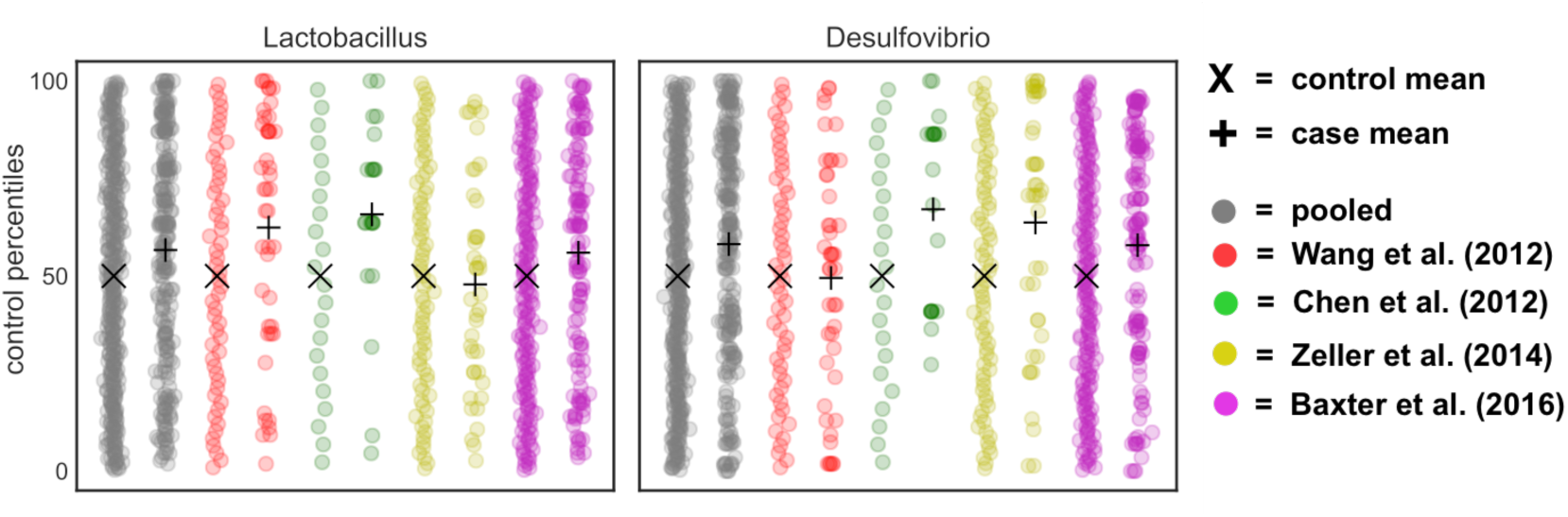
Genera that do not show a significant difference between CRC cases and controls within a given study, but do after pooling. Two genera did not show significant differences between cases and controls within a study, but became significant after pooling across CRC studies (q ≤ 0.05).

While prior work has suggested that there may not be consistent associations between the gut microbiome and obesity [35], we observed six genera in a recent meta-analysis that differed significantly across two or more (out of five) independent obesity studies [17]. Of these six genera, four (*Roseburia*, *Clostridium IV*, *Oscillobacter*, and *Pseudoflavonifractor*) were also found to be significant in the pooled, percentile-normalized results (Table S1; Fig. S2). The two remaining genera not found to be significant in the percentile-normalized analysis (*Mogibacterium* and *Anaerovorax*) showed highly irregular responses across the 11 obesity studies analyzed in this study (Fig. S3). Despite the irregular behavior of these genera, Fisher’s and Stouffer’s methods both identified *Mogibacterium* as significantly associated with obesity (Table S1).

## Discussion

Batch effects are unavoidable when working with high-throughput data generation platforms. The RNA microarray community has been proactive in the development of tools for dealing with these effects [1, 8]. However, these tools are not as effective when batch effects are confounded with biological signals or when parametric assumptions do not apply, which is often the case in microbiome case-control studies. Therefore, model-free methods are needed for correcting batch effects across microbiome datasets. Fortunately, case-control studies can be internally normalized by their own control samples. Any study-specific batch effects in the case samples will be present in the control samples, and by converting the case data into percentiles of the control distribution these effects are attenuated without making parametric assumptions.

Relative abundance, limma-corrected, and ComBat-corrected data - but not percentile-normalized data - quickly yielded a large number of spurious results when cases from one study were tested against controls from another (Fig. 3). Additionally, when control populations from different batches were compared to one another, non-normalized data yielded a much larger number of false positives than batch-corrected data (Fig. 4). Our percentile-normalization approach was much more effective than limma and ComBat in controlling false positives (Fig. 4), especially in the presence of low-abundance taxa (Fig. S1).

Because pooling datasets increases statistical power, it is tempting to pool these data even in the absence of suitable batch-correction methods. Consequently, pooling non-normalized data from different batches has been common practice in the microbiome field [27, 29, 35, 47–55]. In this paper, we demonstrate why this practice is highly inadvisable. Pooling batch-corrected data from multiple studies allowed us to detect significant differences that were not found within a given study (Figs. 5 and 7), while removing associations that were weak or inconsistent across studies (Fig. 6 and S3). Percentile-normalized results often identified significant differences between cases and controls that were missed by other normalization methods (Table 1). For CDI, percentile-normalized results identified about the same number of significant hits as the other batch-correction methods (Table 1), which was likely due to the fact that the biological signal associated with diarrhea is very strong [17]. In cases where the biological signal is strong, results should be robust to the types of analyses employed. For UC and CD studies (IBD), percentile-normalization identified several significant genera that limma and ComBat did not (Table 1). The reduced number of significant hits from limma- and ComBat-corrected data for IBD was likely due to heterogeneous control cohorts across these studies (i.e. healthy patients vs. non-IBD patients), which likely smoothed-out inflammation-associated signals. This result highlights the importance of having consistent definitions for case and control cohorts across studies.

We compared percentile-normalization and pooling to Fisher’s and Stouffer’s methods for combining independent p-values. Stouffer’s method is similar to Fisher’s, but includes weights for each p-value based on the number of samples in a study. Percentile-normalization consistently identified a larger number of significant hits than Stouffer’s and Fisher’s methods, confirming that pooling data increases sensitivity (i.e. reducing putative false negatives). Methods for combining p-values from independent studies are quite robust and should probably be considered as a safe alternative to pooling (i.e. lower chance of false positives). However, in the case of our obesity analysis, Fisher’s and Stouffer’s methods identified *Mogibacterium* as significant despite its apparent inconsistency across studies (Fig. S3).

In conclusion, we present a robust, model-free procedure for transforming each feature in a microbiome case-control dataset into percentiles of its control distribution (Fig. 1). The main conditions for applying this method are that 1) each batch must have a sizeable number of control samples (i.e. the density of the control distribution limits the resolution of the percentile-transformation of the case samples), and 2) case and control populations should be consistently defined across batches (i.e. same definition of ‘healthy’ or ‘diseased’ groups). Given these caveats, percentile-normalized features can be pooled across studies for univariate statistical testing (whichever test a researcher prefers - ideally non-parametric), alleviating the batch effect problem. This model-free procedure could also be applied to other types of ‘omics datasets with consistently defined internal controls. We find that this procedure allows us to identify differences between cases and controls that are often missed by more conservative meta-analysis techniques. Methods developed for batch-correction in microarray data, like limma and ComBat, can partially reduce batch effects in microbiome studies (Figs. 2-4), but appear to obscure real patterns if batch effects are not independent of biological signals or if the parametric assumptions of these models are not valid. We suggest that methods like limma and ComBat are useful for studies lacking case and control groups. However, when studies have consistently defined internal controls, percentile-normalization should be the preferred batch correction approach. Future work should focus on developing parametric models specifically for batch correction in microbiome datasets, which could further improve sensitivity to detect subtle biological differences across studies.

## Acknowledgements

We thank Sean Kearney and members of the Alm and Irizarry labs for helpful feedback and Marc Sze and Patrick Schloss for providing online code to download the Wu et al. (2011) and Zeevi et al. (2015) obesity datasets.

## Supplemental Materials

**Table S1.** Excel file containing tabs for each disease analyzed in this study: each tab contains information on significant genera from Table 1.

**Figure S1.**
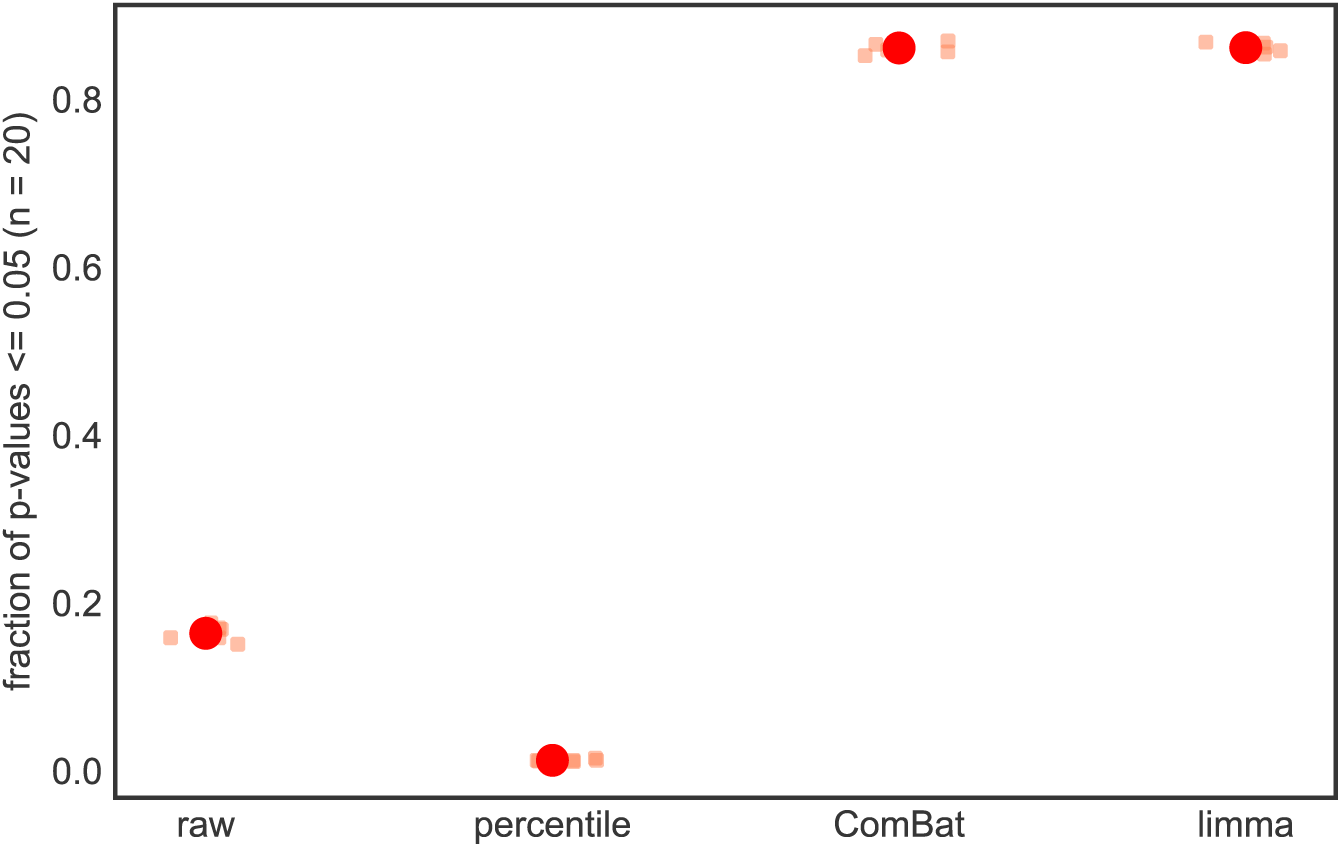
False positive rates are reduced by percentile-normalization, but not by ComBat or limma, in the presence of low-abundance OTUs. Random sets of 40 Baxter controls and random sets of 40 Zeller controls were selected for null case-control comparisons (20 iterations). Smaller points show the fraction of p-values ≤ 0.05 within a given iteration, while larger dots show the average value across all 20 iterations. Within each category, smaller points are randomly jittered along the x-axis for better visualization. The fraction of p-values ≤ 0.05 is highly inflated for all methods except percentile-normalization (red dashed line shows the null-expectation for p-values). All OTUs were included in this analysis (i.e. no abundance filter prior to running tests). ComBat and limma show highly skewed p-value distributions when including low-abundance OTUs.

**Fig. S2.**
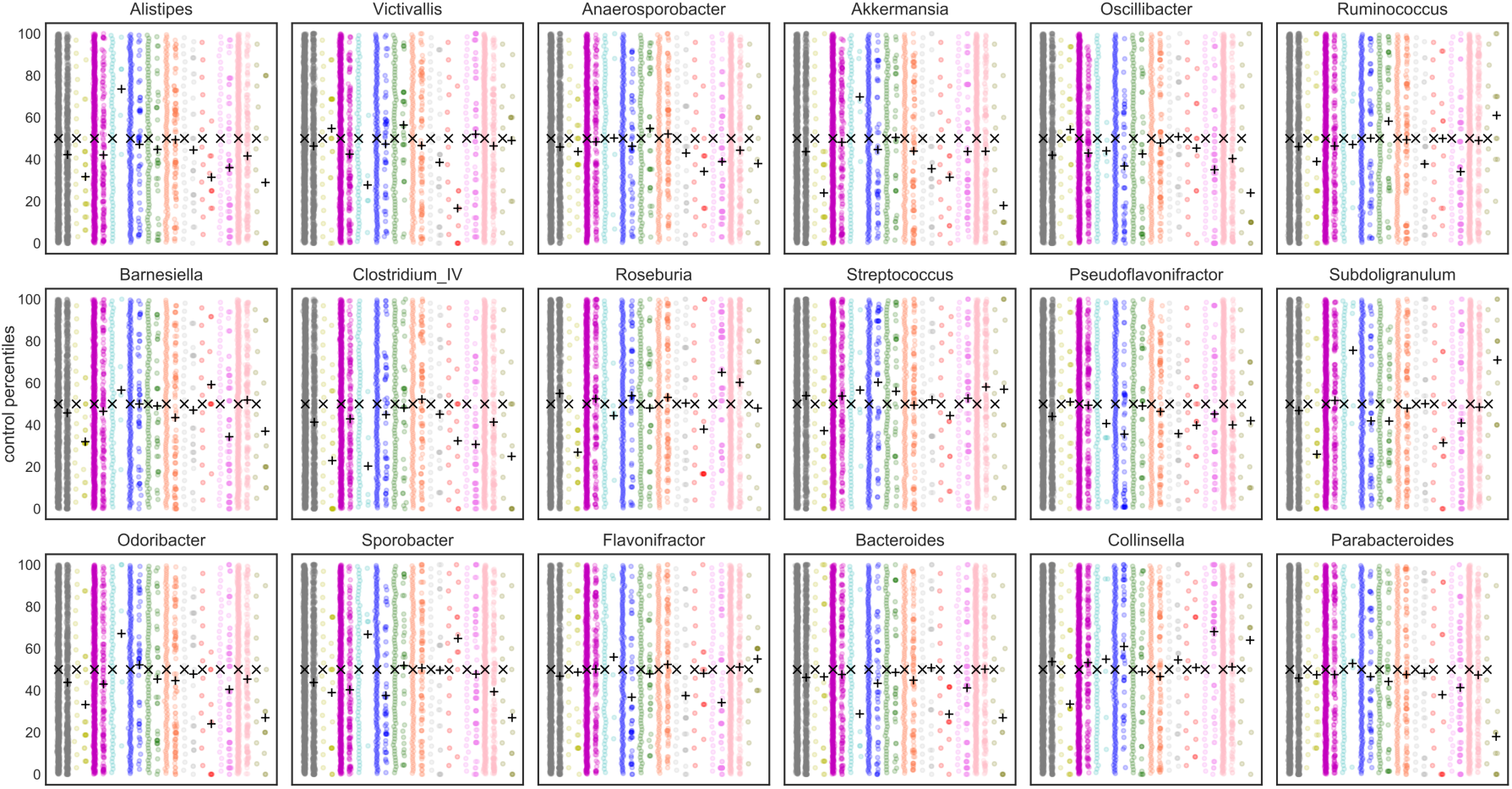
18 genera significantly different (q ≤ 0.05) between obese and healthy patients based on pooled percentile-normalized analysis (from Table 1) Gray points show data pooled from all 11 obesity studies. Other colors show individual obesity studies. Studies listed in order from left to right: Zhu et al. (2014), Zeevi et al. (2015), Wu et al. (2011), Baxter et al. (2016), Schubert et al. (2014), Zupancic et al. (2012), Ross et al. (2015), Jumpertz et al. (2011), Turnbaugh et al. (2009), Goodrich et al. (2014), Escobar et al. (2014). X symbols indicate the average control percentiles, while + symbols indicate the average case percentiles.

**Fig. S3.**
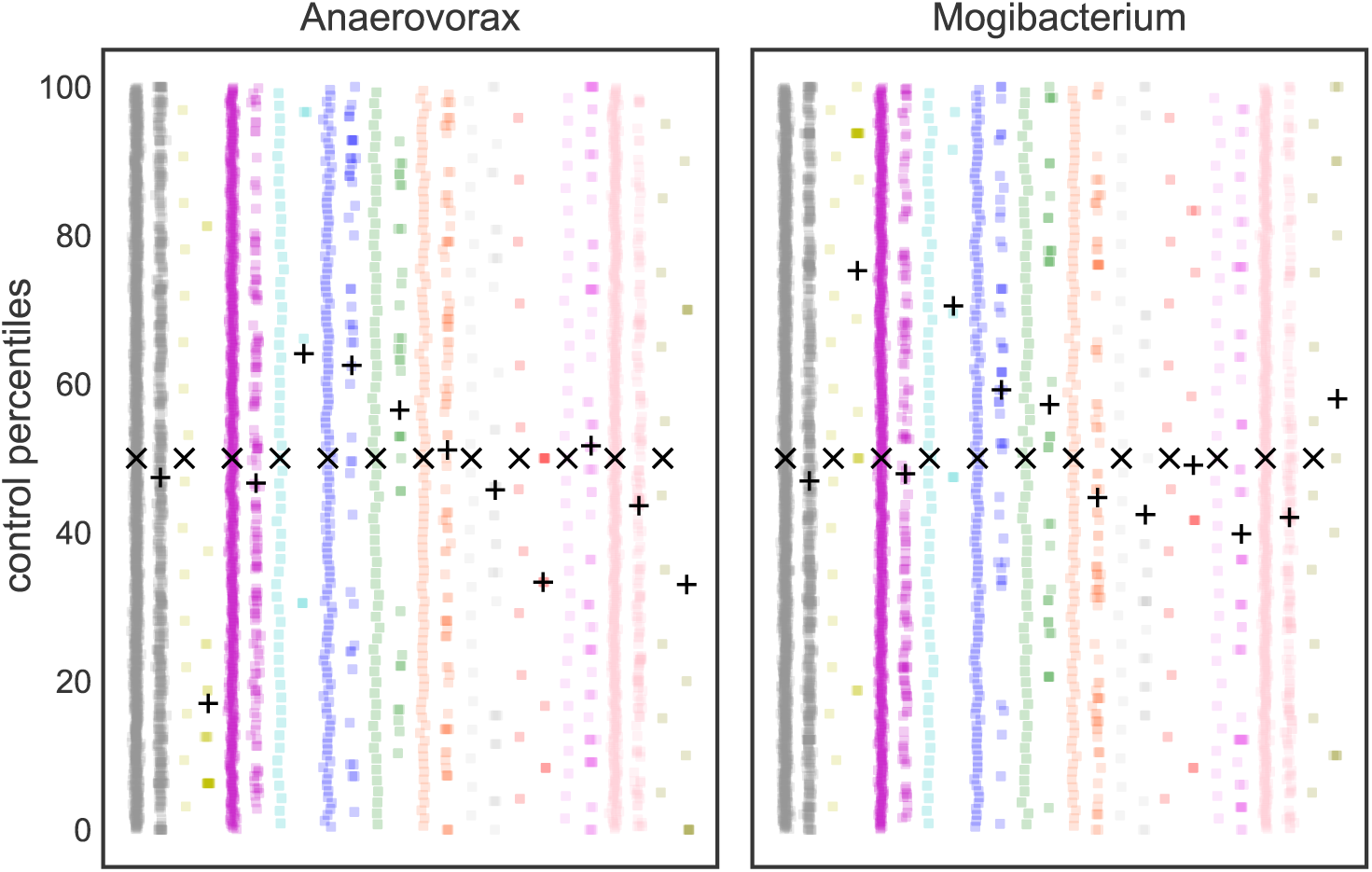
Two genera that were significant (q ≤ 0.05) in at least two obesity studies in Duvallet et al. (2017), but were not significant in the pooled, percentile-normalized analysis. Gray points show data pooled from all 11 obesity studies. Other colors show individual obesity studies. Studies listed in order from left to right: Zhu et al. (2014), Zeevi et al. (2015), Wu et al. (2011), Baxter et al. (2016), Schubert et al. (2014), Zupancic et al. (2012), Ross et al. (2015), Jumpertz et al. (2011), Turnbaugh et al. (2009), Goodrich et al. (2014), Escobar et al. (2014). X symbols indicate the average control percentiles, while + symbols indicate the average case percentiles.

## References

1. Leek JT, Scharpf RB, Bravo HC, Simcha D, Langmead B, Johnson WE, et al. Tackling the widespread and critical impact of batch effects in high-throughput data. Nat Rev Genet. 2010;11(10):733–9.

2. Goh WWB, Wang W, Wong L. Why batch effects matter in omics data, and how to avoid them. Trends Biotechnol. 2017;35(6):498–507.

3. Schloss PD, Gevers D, Westcott SL. Reducing the Effects of PCR Amplification and Sequencing Artifacts on 16S rRNA-Based Studies. PLoS One. 2011;6(12):e27310. doi: 10.1371/journal.pone.0027310.

4. Leek JT, Storey JD. Capturing heterogeneity in gene expression studies by surrogate variable analysis. PLoS Genet. 2007;3(9):e161.

5. Ritchie ME, Phipson B, Wu D, Hu Y, Law CW, Shi W, et al. limma powers differential expression analyses for RNA-sequencing and microarray studies. Nucleic Acids Res. 2015;43(7):e47-e.

6. Alter O, Brown PO, Botstein D. Singular value decomposition for genome-wide expression data processing and modeling. Proc Natl Acad Sci USA. 2000;97(18):10101–6.

7. Benito M, Parker J, Du Q, Wu J, Xiang D, Perou CM, et al. Adjustment of systematic microarray data biases. Bioinformatics. 2004;20(1):105–14.

8. Chen C, Grennan K, Badner J, Zhang D, Gershon E, Jin L, et al. Removing Batch Effects in Analysis of Expression Microarray Data: An Evaluation of Six Batch Adjustment Methods. PLoS One. 2011;6(2):e17238. doi: 10.1371/journal.pone.0017238.

9. Johnson WE, Li C, Rabinovic A. Adjusting batch effects in microarray expression data using empirical Bayes methods. Biostatistics. 2007;8(1):118–27.

10. Weiss S, Amir A, Hyde ER, Metcalf JL, Song SJ, Knight R. Tracking down the sources of experimental contamination in microbiome studies. Genome Biol. 2014;15(12):564. doi: 10.1186/s13059-014-0564-2.

11. Shen H, Rogelj S, Kieft TL. Sensitive, real-time PCR detects low-levels of contamination by Legionella pneumophila in commercial reagents. Mol Cell Probes. 2006;20. doi: 10.1016/j.mcp.2005.09.007.

12. Nguyen NH, Smith D, Peay K, Kennedy P. Parsing ecological signal from noise in next generation amplicon sequencing. New Phytol. 2015;205(4):1389–93. doi: 10.1111/nph.12923.

13. Gibbons SM. The Built Environment Is a Microbial Wasteland. mSystems. 2016;1(2):e00033–16.

14. Chase J, Fouquier J, Zare M, Sonderegger DL, Knight R, Kelley ST, et al. Geography and Location Are the Primary Drivers of Office Microbiome Composition. mSystems. 2016;1(2). doi: 10.1128/mSystems.00022-16.

15. Fisher RA. Statistical methods for research workers: Genesis Publishing Pvt Ltd; 1925.

16. Stouffer SA. Adjustment during army life: Princeton University Press; 1949.

17. Duvallet C, Gibbons SM, Gurry T, Irizarry RA, Alm EJ. Meta-analysis of gut microbiome studies identifies disease-specific and shared responses. Nat Comm. 2017;8(1):1784.

18. Baxter NT, Ruffin MT, Rogers MA, Schloss PD. Microbiota-based model improves the sensitivity of fecal immunochemical test for detecting colonic lesions. Genom Med. 2016;8(1):37.

19. Zeller G, Tap J, Voigt AY, Sunagawa S, Kultima JR, Costea PI, et al. Potential of fecal microbiota for early-stage detection of colorectal cancer. Mol Sys Biol. 2014;10(11):766.

20. Zackular JP, Rogers MA, Ruffin MT, Schloss PD. The human gut microbiome as a screening tool for colorectal cancer. Cancer Prev Res. 2014;7(11):1112–21.

21. Chen W, Liu F, Ling Z, Tong X, Xiang C. Human intestinal lumen and mucosa-associated microbiota in patients with colorectal cancer. PLoS One. 2012;7(6):e39743.

22. Gevers D, Kugathasan S, Denson LA, Vázquez-Baeza Y, Van Treuren W, Ren B, et al. The treatment-naive microbiome in new-onset Crohn’s disease. Cell Host Microbe. 2014;15(3):382–92.

23. Papa E, Docktor M, Smillie C, Weber S, Preheim SP, Gevers D, et al. Non-invasive mapping of the gastrointestinal microbiota identifies children with inflammatory bowel disease. PLoS One. 2012;7(6):e39242.

24. Morgan XC, Tickle TL, Sokol H, Gevers D, Devaney KL, Ward DV, et al. Dysfunction of the intestinal microbiome in inflammatory bowel disease and treatment. Genome Biol. 2012;13(9):R79.

25. Willing BP, Dicksved J, Halfvarson J, Andersson AF, Lucio M, Zheng Z, et al. A pyrosequencing study in twins shows that gastrointestinal microbial profiles vary with inflammatory bowel disease phenotypes. Gastroenterology. 2010;139(6):1844–54. e1.

26. Turnbaugh PJ, Hamady M, Yatsunenko T, Cantarel BL, Duncan A, Ley RE, et al. A core gut microbiome in obese and lean twins. Nature. 2009;457(7228):480–4.

27. Escobar JS, Klotz B, Valdes BE, Agudelo GM. The gut microbiota of Colombians differs from that of Americans, Europeans and Asians. BMC Microbiol. 2014;14(1):311.

28. Zupancic ML, Cantarel BL, Liu Z, Drabek EF, Ryan KA, Cirimotich S, et al. Analysis of the gut microbiota in the old order Amish and its relation to the metabolic syndrome. PLoS One. 2012;7(8):e43052.

29. Ross MC, Muzny DM, McCormick JB, Gibbs RA, Fisher-Hoch SP, Petrosino JF. 16S gut community of the Cameron County Hispanic Cohort. Microbiome. 2015;3(1):7.

30. Goodrich JK, Waters JL, Poole AC, Sutter JL, Koren O, Blekhman R, et al. Human genetics shape the gut microbiome. Cell. 2014;159(4):789–99.

31. Zeevi D, Korem T, Zmora N, Israeli D, Rothschild D, Weinberger A, et al. Personalized nutrition by prediction of glycemic responses. Cell. 2015;163(5):1079–94.

32. Wu GD, Chen J, Hoffmann C, Bittinger K, Chen Y-Y, Keilbaugh SA, et al. Linking long-term dietary patterns with gut microbial enterotypes. Science. 2011;334(6052):105–8.

33. Schubert AM, Rogers MA, Ring C, Mogle J, Petrosino JP, Young VB, et al. Microbiome data distinguish patients with Clostridium difficile infection and non-C. difficile-associated diarrhea from healthy controls. mBio. 2014;5(3):e01021–14.

34. Jumpertz R, Le DS, Turnbaugh PJ, Trinidad C, Bogardus C, Gordon JI, et al. Energy-balance studies reveal associations between gut microbes, caloric load, and nutrient absorption in humans. Am J Clin Nutr. 2011;94(1):58–65.

35. Sze MA, Schloss PD. Looking for a Signal in the Noise: Revisiting Obesity and the Microbiome. MBio. 2016;7(4):e01018–16.

36. Vincent C, Stephens DA, Loo VG, Edens TJ, Behr MA, Dewar K, et al. Reductions in intestinal Clostridiales precede the development of nosocomial Clostridium difficile infection. Microbiome. 2013;1(1):18.

37. Edgar RC. Search and clustering orders of magnitude faster than BLAST. Bioinformatics. 2010;26(19):2460–1.

38. Wang Q, Garrity GM, Tiedje JM, Cole JR. Naïve Bayesian Classifier for Rapid Assignment of rRNA Sequences into the New Bacterial Taxonomy. Appl Environ Microbiol. 2007;73(16):5261–7. doi: 10.1128/aem.00062-07.

39. Pedregosa F, Varoquaux G, Gramfort A, Michel V, Thirion B, Grisel O, et al. Scikit-learn: Machine learning in Python. J Mach Learning Res. 2011;12(Oct):2825–30.

40. Jones E, Oliphant T, Peterson P. SciPy: Open source scientific tools for Python, 2001–. URL http://www.scipyorg.. 2007;73:86.

41. Seabold S, Perktold J, editors. Statsmodels: Econometric and statistical modeling with python. Proceedings of the 9th Python in Science Conference; 2010.

42. Oksanen J. Multivariate analysis of ecological communities in R: vegan tutorial. R package version. 2011;1(7).

43. Tjalsma H, Boleij A, Marchesi JR, Dutilh BE. A bacterial driver–passenger model for colorectal cancer: beyond the usual suspects. Nat Rev Microbiol. 2012;10(8):575–82.

44. Zhu L, Baker RD, Baker SS. Gut microbiome and nonalcoholic fatty liver diseases. Pediatric Res. 2014;77(1-2):245–51.

45. Wang T, Cai G, Qiu Y, Fei N, Zhang M, Pang X, et al. Structural segregation of gut microbiota between colorectal cancer patients and healthy volunteers. ISME J. 2012;6(2):320–9.

46. Glass GV. Primary, secondary, and meta-analysis of research. Educ Res. 1976;5(10):3–8.

47. Oliveira FS, Brestelli J, Cade S, Zheng J, Iodice J, Fischer S, et al. MicrobiomeDB: a systems biology platform for integrating, mining and analyzing microbiome experiments. Nucleic Acids Res. 2017;46(D1):D684–D91.

48. Huse SM, Dethlefsen L, Huber JA, Welch DM, Relman DA, Sogin ML. Exploring microbial diversity and taxonomy using SSU rRNA hypervariable tag sequencing. PLoS Genet. 2008;4(11):e1000255.

49. Koren O, Knights D, Gonzalez A, Waldron L, Segata N, Knight R, et al. A guide to enterotypes across the human body: meta-analysis of microbial community structures in human microbiome datasets. PLoS Comput Biol. 2013;9(1):e1002863.

50. Krych L, Hansen CH, Hansen AK, Van Den Berg FW, Nielsen DS. Quantitatively different, yet qualitatively alike: a meta-analysis of the mouse core gut microbiome with a view towards the human gut microbiome. PLoS One. 2013;8(5):e62578.

51. Lozupone CA, Li M, Campbell TB, Flores SC, Linderman D, Gebert MJ, et al. Alterations in the gut microbiota associated with HIV-1 infection. Cell Host Microbe. 2013;14(3):329–39.

52. Lozupone CA, Stombaugh J, Gonzalez A, Ackermann G, Wendel D, Vazquez-Baeza Y, et al. Meta-analyses of studies of the human microbiota. Genome Res. 2013;23. doi: 10.1101/gr.151803.112.

53. Walters WA, Xu Z, Knight R. Meta-analyses of human gut microbes associated with obesity and IBD. FEBS Lett. 2014;588(22):4223–33.

54. Pasolli E, Truong DT, Malik F, Waldron L, Segata N. Machine learning meta-analysis of large metagenomic datasets: tools and biological insights. PLoS Comput Biol. 2016;12(7):e1004977.

55. Vogtmann E, Hua X, Zeller G, Sunagawa S, Voigt AY, Hercog R, et al. Colorectal cancer and the human gut microbiome: reproducibility with whole-genome shotgun sequencing. PLoS One. 2016;11(5):e0155362.

